# The WAVE complex drives the morphogenesis of the photoreceptor outer segment cilium

**DOI:** 10.1101/2022.11.21.517374

**Authors:** William J. Spencer, Nicholas F. Schneider, Nikolai P. Skiba, Vadim Y. Arshavsky

**Affiliations:** Department of Ophthalmology, Duke University Medical Center, Durham, NC 27710; Department of Pharmacology and Cancer Biology, Duke University Medical Center, Durham, NC 27710

**Keywords:** photoreceptor, outer segment, actin cytoskeleton, WAVE, SCAR, Wasf3, Arp2/3, cilium

## Abstract

The photoreceptor outer segment is a modified cilium filled with hundreds of flattened “disc” membranes responsible for efficient light capture. To maintain photoreceptor health and functionality, outer segments are continuously renewed through the addition of new discs at their base. This process is driven by branched actin polymerization nucleated by the Arp2/3 complex. To induce actin polymerization, Arp2/3 requires a nucleation promoting factor. Here, we show that the nucleation promoting factor driving disc morphogenesis is the pentameric WAVE complex and identify all protein subunits of this complex. We further demonstrate that the knockout of one of them, WASF3, abolishes actin polymerization at the site of disc morphogenesis leading to formation of disorganized membrane lamellae emanating from the photoreceptor cilium instead of an outer segment. These data establish that, despite the intrinsic ability of photoreceptor ciliary membranes to form lamellar structures, WAVE-dependent actin polymerization is essential for organizing these membranes into a proper outer segment.

## Introduction

The light-sensitive membranes of vertebrate photoreceptors are flattened membrane vesicles, called “discs”, which are stacked by the hundreds inside the photoreceptor cell’s ciliary outer segment organelle. Outer segments are continuously renewed by the daily addition of new discs at their base and phagocytosis of old discs from their tips by the retinal pigment epithelium (see (1) for a recent review). This renewal process is thought to play a critical role in maintaining photoreceptor health by recycling the material damaged upon continuous light exposure. Disc formation begins with the evagination of the ciliary membrane at the base of the outer segment, which is subsequently flattened into a lamella, expanded to the diameter of the outer segment and, at least in the case of rod photoreceptors, is enclosed inside the outer segment (2-5). Ciliary membrane evagination is driven by the filamentous actin (F-actin) network expanding at this site (6,7). Recently, we showed that this network is branched and assembled by the actin nucleating complex Arp2/3 (8). These findings suggest that disc formation is mediated by a cycle of branched actin polymerization and depolymerization, repeatedly pushing out the ciliary plasma membrane to create each new disc. This mechanism is analogous to the action of lamellipodia at the leading edge of migrating cells, which are dynamic actin-driven membrane protrusions nucleated by Arp2/3 (9). In this mechanism, two subunits of the Arp2/3 complex, acting as structural homologs of actin, associate with an actin monomer thereby mimicking the structure of an actin trimer. This conformation induces the spontaneous addition of actin monomers leading to filament elongation and F-actin network expansion (10-13).

However, Arp2/3 alone is insufficient to initiate actin polymerization. This is because Arp2/3’s binding to an actin monomer requires the monomer to first associate with one of the nucleation promoting factors (14-16), such as in the example illustrated schematically in Fig. 1. Nucleation promoting factors are essential not only for Arp2/3-mediated actin nucleation but also regulation of Arp2/3 activity, both temporally and spatially (15,16). This regulation is conferred through the diversity of the nucleation promoting factors and their interacting partners. For example, the monomeric WASP protein is engaged in Arp2/3-dependent endocytosis and phagocytosis, whereas the heteropentameric WAVE complex drives lamellipodia formation (17). Further enhancing the functional specialization of branched actin networks, there are multiple protein isoforms of both WASP and the individual subunits of the WAVE complex (17,18).

**Figure 1.**
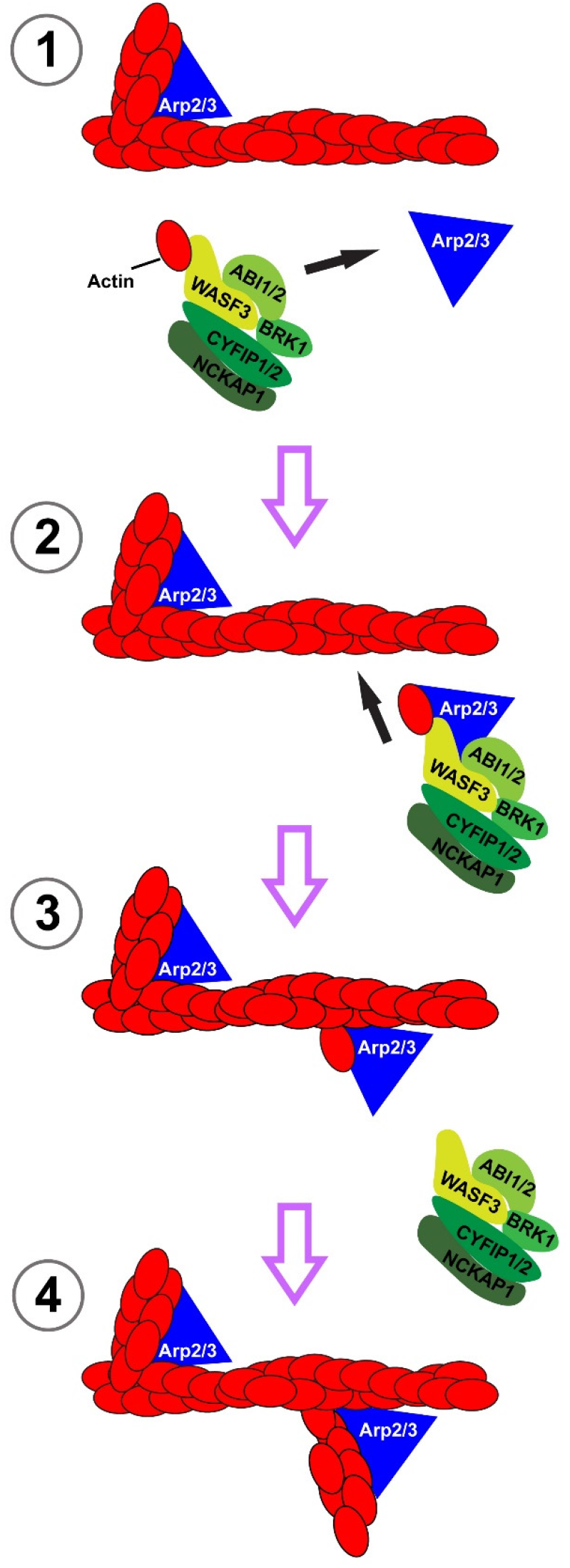
The roles of the Arp2/3 and WAVE complexes in building branched actin networks. Arp2/3 is labeled in blue, WAVE in green and actin in red. *In step 1*, the WAVE complex whose WASF subunit is bound to an actin monomer interacts with Arp2/3. *In step 2*, Arp2/3 associated with both monomeric actin and the WAVE complex binds to the side of a preexisting actin filament. *In step 3*, the WAVE complex disassociates from the filament leaving Arp2/3 behind. *In step 4*, the newly seeded actin branch elongates. See (17) for a review. The indicated WAVE complex subunits represent specific isoforms identified in this study.

In this study, we have found that the only nucleation promoting factor for Arp2/3 in the outer segment is the WAVE complex. We identified the protein isoforms representing all five subunits of this complex as WASF3, ABI1/2, NCKAP1, CYFIP1/2 and BRK1. We next showed that the knockout of WASF3 – a protein directly interacting with Arp2/3 – causes a specific loss of F-actin at the site of disc morphogenesis and abolishes the formation of the outer segment. In place of outer segments, *Wasf3*^*-/-*^ photoreceptors elaborate large membrane whorls emanating from their cilia. These whorls are composed of discrete membrane layers, suggesting that photoreceptor ciliary membranes have an intrinsic capacity to self-organize into lamellae even without actin polymerization. These data demonstrate that the functional specialization of the Arp2/3 complex in outer segment morphogenesis is conferred by the WAVE complex containing WASF3. Without WAVE-Arp2/3 driven actin polymerization, photoreceptor ciliary membranes retain the ability to flatten but fail to form well-organized disc stacks.

## Results

### The WAVE complex is the major nucleation promoting factor in photoreceptor outer segments

To identify the nucleation promoting factor(s) working with Arp2/3 at the site of disc morphogenesis, we conducted proteomic analysis of rod outer segments isolated from bovine retinas. Among all proteins confidently identified in this proteome (see Methods for protein identification criteria), the only known nucleation promoting factor was WASF3, along with two other components of the pentameric WAVE complex, CYFIP2 and NCKAP1 (Supplementary Table 1). Consistently, WASF3 was detected in another, related proteome of membrane vesicles released from the photoreceptor cilia of *rds* mice, which are enriched in proteins normally involved in disc formation, including actin and Arp2/3 (8). The hypothesis that WASF3 acts as a nucleation promoting factor for Arp2/3 at the disc morphogenesis site was put forward in a recent study (19). Using yeast two-hybrid system and protein expression in cell culture, the authors showed that WASF3 interacts with PCARE, a protein whose knockout was previously shown to cause severe disruption of outer segment membranes (20). They further showed that these proteins co-localize at the site of disc formation and that PCARE knockout eliminates both the F-actin patch and WASF3 at this site. Finally, they showed that PCARE recruits WASF3 to the cilium when these proteins are ectopically co-expressed in cell culture, which leads to the expansion of ciliary F-actin. With strong evidence in place, our goal was to directly test whether WASF3 is an indispensable regulator of the actin network driving photoreceptor disc formation.

We first identified all members of the WAVE complex present in photoreceptor outer segments. This was accomplished by immunoprecipitating WASF3 from a lysate of purified mouse rod outer segments and identifying co-precipitating proteins by mass spectrometry. We found that the immunoprecipitate contained the WAVE complex proteins WASF3, ABI1, ABI2, CYFIP1, CYFIP2 and NCKAP1 (Fig. 2A and Supplementary Table 2). To confirm this result and to determine whether the other WASF isoforms, WASF1 or WASF2, may also be present in the outer segment, we conducted a reciprocal precipitation of the WAVE complex using an anti-ABI1 antibody. We identified WASF3, CYFIP1, CYFIP2 and NCKAP1 co-precipitating with ABI1 (Fig 2A and Supplementary Table 3). These data pinpoint WASF3 as a potentially major player in nucleating actin at the site of the disc morphogenesis. This hypothesis is particularly intriguing because WASF3 has been found to contribute to metastatic invasion of cancer cells but has yet to be directly shown to play a critical role in any normal biological function (21,22).

**Figure 2.**
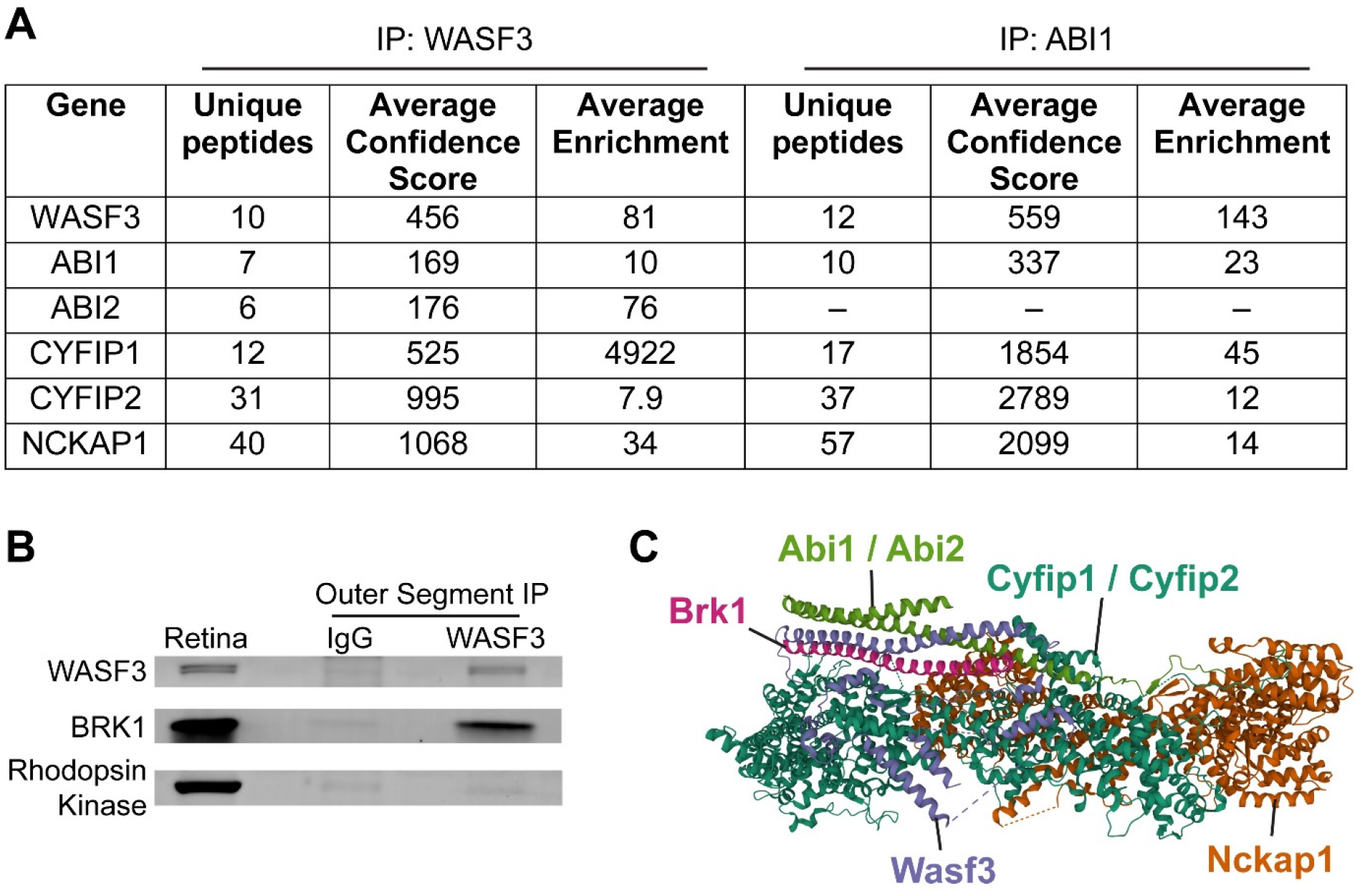
The pentameric WAVE complex is the nucleation promoting factor in photoreceptor outer segments. (A) Specific protein isoforms of the WAVE complex identified by mass spectrometry in immunoprecipitates from lysed purified mouse rod outer segments using either anti-WASF3 or anti-ABI1 antibodies. Proteins included in the table had at least a two-fold enrichment over control immunoprecipitates with anti-IgG antibodies. See Supplementary Table 2 for the full list of identified proteins in each experiment. (B) Western blot showing co-precipitation of BRK1 with WASF3 from lysed rod outer segments using the anti-WASF3 antibody. Immunoblotting for rhodopsin kinase, an outer segment protein not associated with the WAVE complex, was used as a negative control. A whole retina lysate was used as input. (C) A ribbon diagram of the WAVE complex labeled with the protein isoforms identified in the photoreceptor outer segment. The structure of the WAVE complex was determined by X-ray crystallography (61) and the ribbon diagram downloaded from World Wide Protein Data Bank (DOI: 10.2210/pdb3P8C/pdb).

The only obligatory component of the WAVE complex missing from our proteomes was BRK1, an 8 kDa protein whose relatively few tryptic peptides may have eluded detection by mass spectrometry. To account for this possibility, we assayed the presence of BRK1 in the outer segment immunoprecipitate obtained with the anti-WASF3 antibody using western blotting and found that BRK1 does co-precipitate with WASF3 (Fig. 2B). Collectively, our data indicate that the WAVE complex in photoreceptor outer segments consists of WASF3, ABI1/ABI2, BRK1, CYFIP1/CYFIP2 and NCKAP1 (Fig. 2C).

### The WAVE complex genes, including *Wasf3*, are robustly expressed in photoreceptors

Complementary evidence suggesting that the WAVE complex containing WASF3 is the predominant nucleation promoting factor in photoreceptors could be found in the previously published RNA-sequencing database from WT rod photoreceptors isolated at postnatal days 2, 4, 6, 10, 14 and 28 (23). These time points span the development of the outer segment, which begins around P7 and continues to ∼P23 (24). The mRNA expression level of *Wasf3* is the highest among all known nucleation promoting factors (Fig. 3A). Furthermore, its expression level increases ∼5-fold during outer segment development in a pattern similar to that of several representative outer segment proteins (Fig. 3B). The expression of *Abi1/2, Cyfip1/2, Nckap1* and *Brk1* was high, although below that of *Wasf3*. Curiously, *Cyfip2* was the only WAVE protein whose temporal pattern of expression correlated with that of *Wasf3*. Considering the remaining known isoforms of the WAVE complex subunits, *Nckap1L* was essentially not expressed, the expression of *Abi3* and *Wasf2* completely diminished by P21, and the expression of *Wasf1* remained very low at all time points (Fig. 3A). Overall, these data are in a strong agreement with our proteomic analysis.

**Figure 3.**
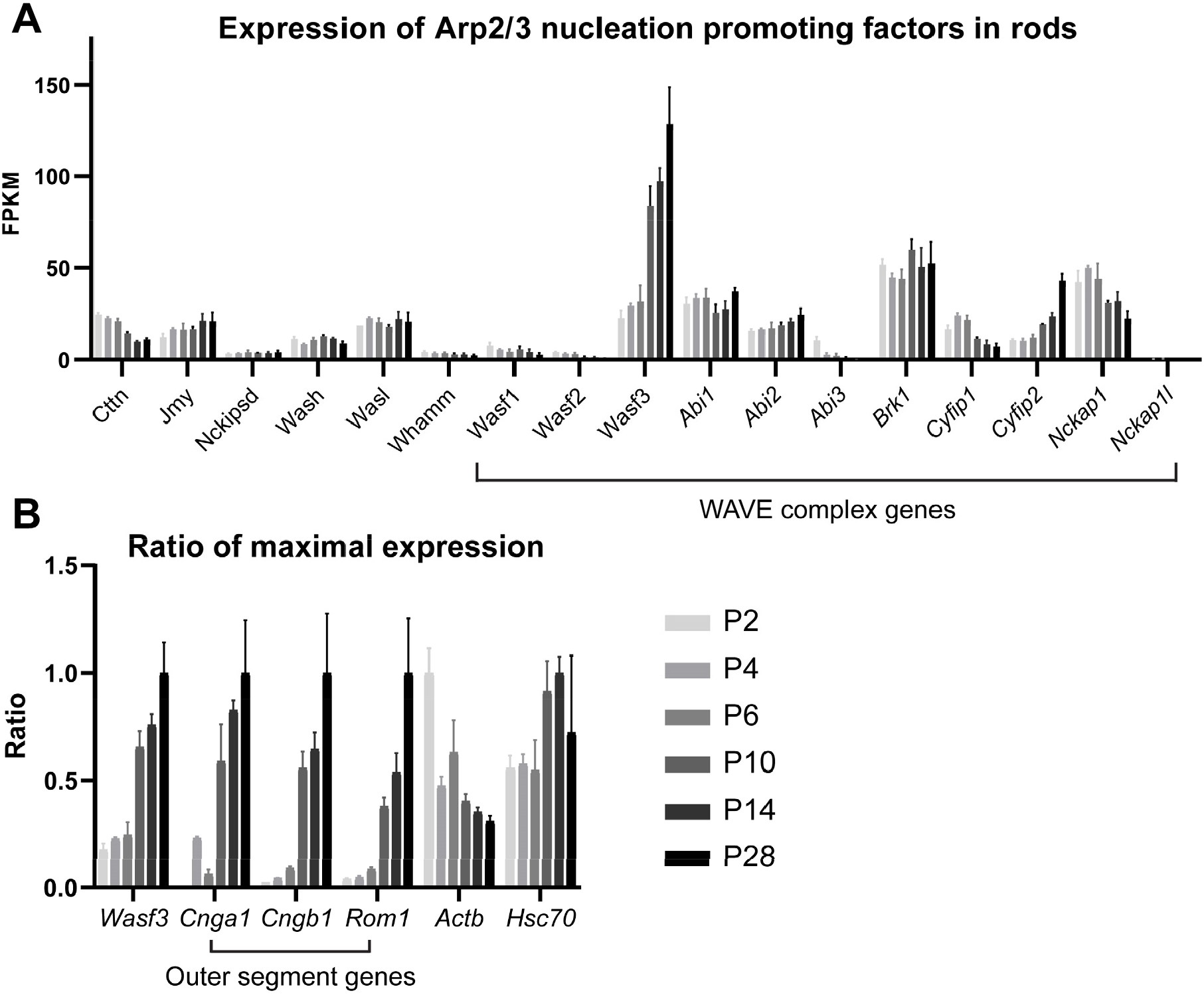
Subunits of the WAVE complex, including WASF3, are highly expressed in photoreceptor cells. (A) mRNA levels of all known nucleation promoting factors expressed in mouse photoreceptor cells. The values are quantified as fragments per kilobase million (FPKM). The data were obtained from a published RNAseq dataset of flow-sorted rod cells isolated at various time points between P2 and P28 (23). Raw data were acquired from NCBI Gene Expression Omnibus series GSE74660. (B) A comparison of the mRNA expression profiles for *Wasf3*, three outer segment proteins (*Cnga1, Cngb1* and *Rom1*), actin (*Actb*) and a housekeeping gene (*Hsc70*). For each gene, the data were normalized to their maximal expression level at any time point. The data in both panels are shown as mean ± SD.

### *Wasf3* ^-/-^ photoreceptors lack F-actin at the site of disc formation

To directly test the role of the WAVE complex containing WASF3 in actin nucleation at the site of disc morphogenesis, we analyzed the retinal phenotype of the *Wasf3* ^*-/-*^ mouse. A previous study of this mouse found no overt morphological or behavioral abnormalities, although the retina was not investigated (25). We first confirmed a complete absence of WASF3 in *Wasf3* ^*-/-*^ retinas by western botting (Supplementary Fig. 1A). We also probed for other WAVE proteins because knockdowns or knockouts of individual subunits typically result in the instability and degradation of the entire complex (26-33). We found that, in the absence of WASF3, all other WAVE proteins were reduced by 24-39%, while the level of the Arp2/3 complex, measured by the abundance of its ARP3 subunit, was unaffected (Supplementary Fig. 1A). We also noted that, in WT retinas, WASF3 immunoblots display a double band, a phenomenon often associated with partial protein phosphorylation. Indeed, the upper band was removed upon treating the sample with phosphatase, consistent with a fraction of WASF3 being phosphorylated in the retina (Supplementary Fig. 1B). This observation is potentially interesting because phosphorylation of WASF3 was shown to regulate its function in other systems (21,34).

The complete loss of WASF3 in *Wasf3* ^*-/-*^ retinas was additionally corroborated by immunostaining retinal sections from WT and knockout mice with an anti-WASF3 antibody. WASF3 staining present throughout all retinal layers of WT mice was absent in the knockout (Supplementary Fig. 1C). In WT photoreceptor cells, WASF3 was distributed throughout the cell body, consistent with WAVE being a soluble protein complex. Confirming a previous report (19), higher magnification images at the base of the outer segment showed that WASF3 staining appears as distinct puncta overlapping with the F-actin staining located at site of disc formation (Fig. 4A).

**Figure 4.**
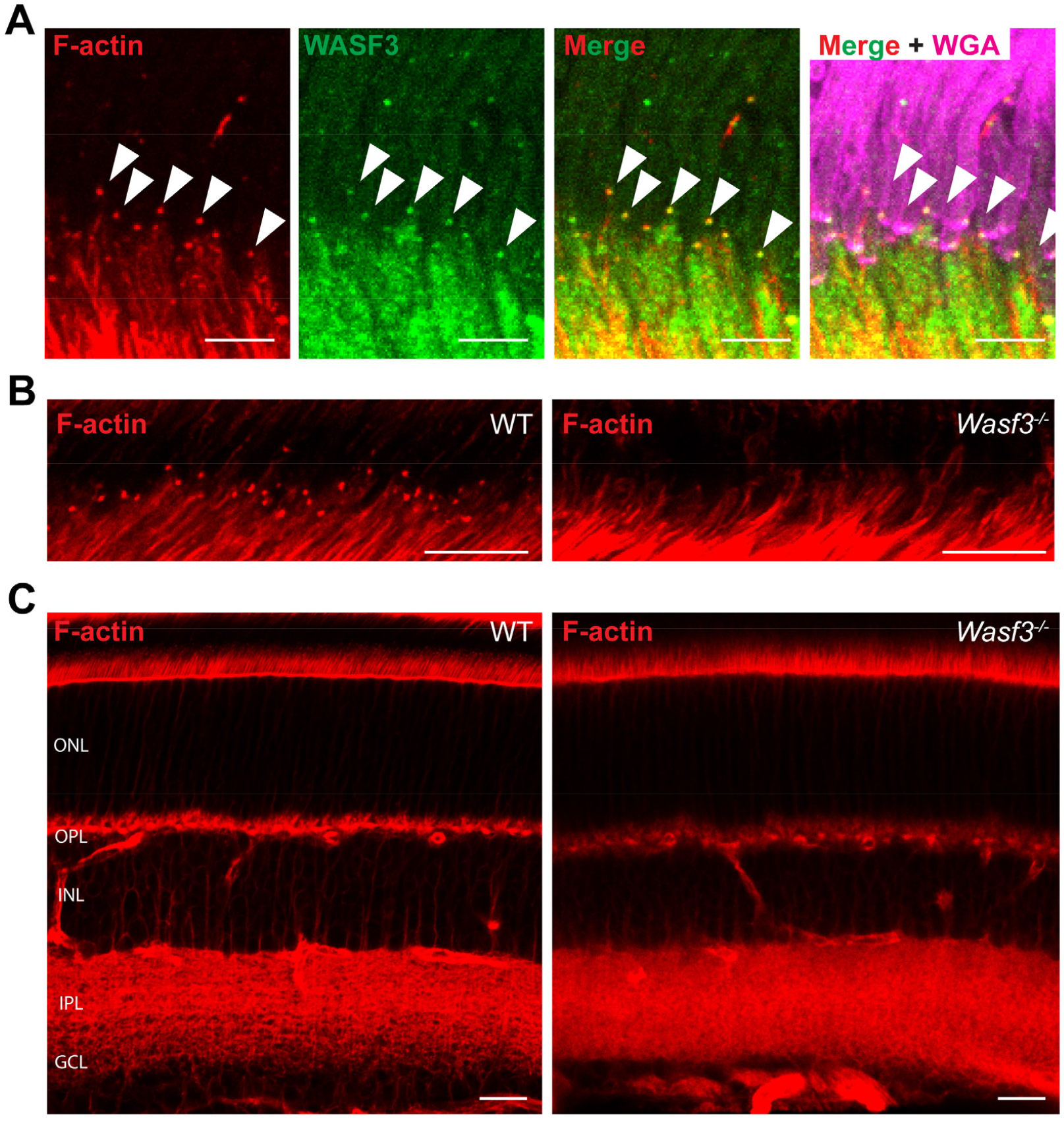
F-actin is specifically lost from the site of disc formation in *Wasf3*^*-/-*^ mice. (A) A retinal cross section, centered at the site of disc formation, from a P21 WT mouse immunostained with anti-WASF3 antibody (green). The sections were co-stained with phalloidin to label F-actin (red) and WGA to label outer segments (magenta). Arrowheads denote F-actin puncta present at the site of disc formation that overlap with distinct puncta of WASF3. Scale bars are 5 μm. (B) Retinal cross sections, centered at the site of disc formation, of P21 WT and *Wasf3* ^*-/-*^ mice stained with phalloidin to label F-actin (red). Scale bars are 10 μm. (C) Whole retinal cross sections of P21 WT and *Wasf3* ^*-/-*^ mice stained with phalloidin. The outer nuclear layer (ONL), outer plexiform layer (OPL), inner nuclear layer (INL), inner plexiform layer (IPL) and ganglion cell layer (GCL) are labeled. The scale bars are 20 μm. See also Supplementary movies 1 and 2.

A previous study showed that the knockout of Arp2/3 from photoreceptor cells results in complete loss of the F-actin network at the site of disc formation (8). To determine whether this were also the case in *Wasf3* ^*-/-*^ mice, we labeled actin filaments in *Wasf3* ^*-/-*^ retinas with fluorophore-conjugated phalloidin and found that the actin network is completely absent at this site (Fig. 4B, Supplementary movies 1 and 2). This loss of F-actin appears to be specific to the outer segment base because F-actin staining elsewhere in the *Wasf3* ^*-/-*^ retina appears normal (Fig. 4C). These results suggest that the WAVE complex containing WASF3 serves as the nucleation promoting factor for Arp2/3 at the site of disc morphogenesis.

In another set of experiments, we considered whether the knockout of WASF3 may affect protein trafficking in photoreceptors, as branched actin polymerization participates in vesicular trafficking in other cell types (35). We analyzed the localization of several representative outer segment proteins in *Wasf3* ^*-/-*^ mice and found that all of them were localized normally, indicating that protein trafficking to the outer segment is unaffected by the loss of WASF3 (Fig. 5A,B). This finding is consistent with our previous observation that the loss of Arp2/3 in rods does not affect protein trafficking to the outer segment (8).

**Figure 5.**
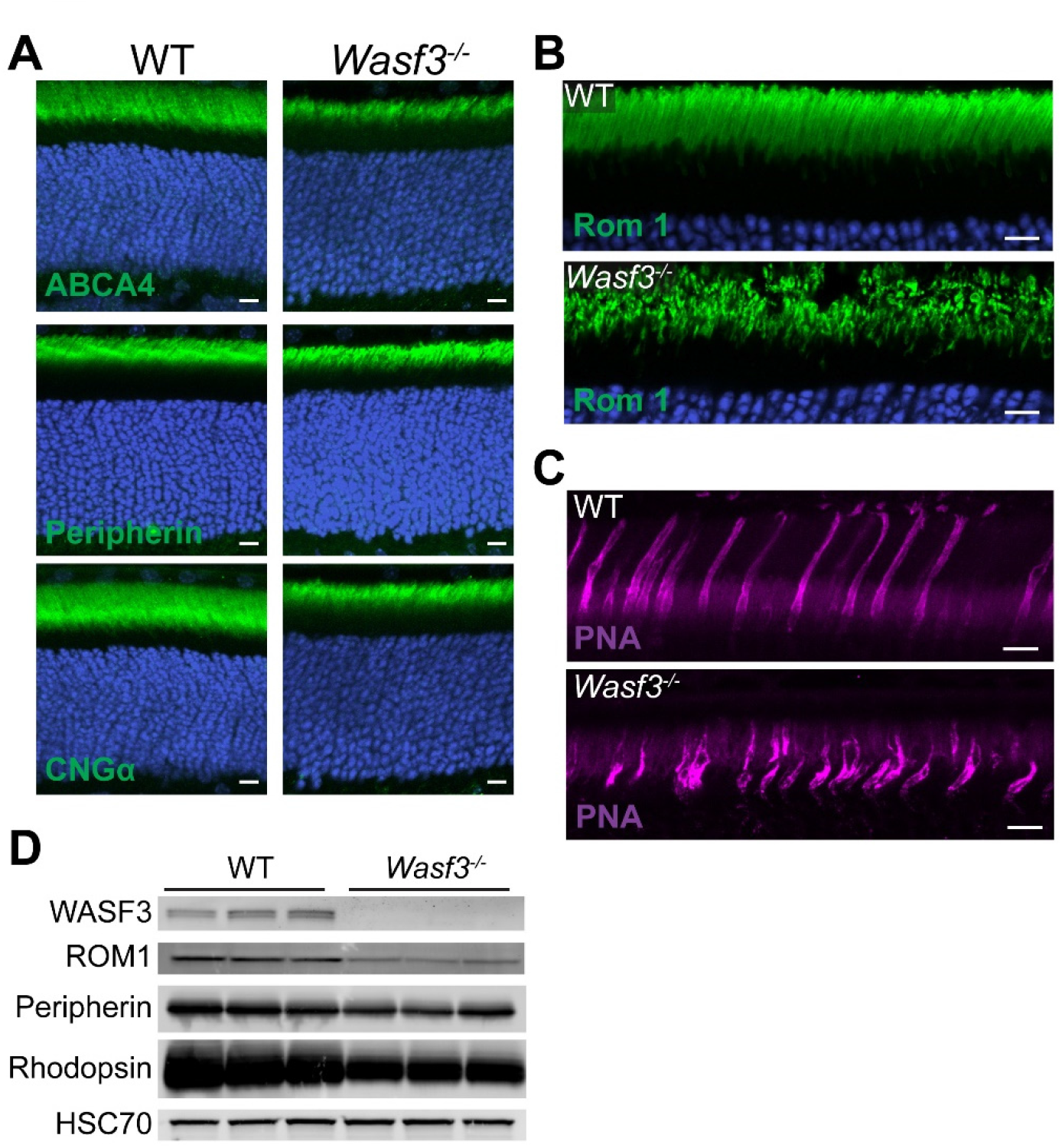
WASF3 knockout results in normal protein trafficking but disrupted outer segment morphology. (A) Retinal cross sections of P21 WT and *Wasf3* ^*-/-*^ mice stained with antibodies against indicated outer segment proteins (green). Nuclei were stained with Hoechst (blue). (B) Expanded view of the outer segment layer in retinal cross sections of P21 WT and *Wasf3* ^*-/-*^ mice stained with anti-ROM1 antibodies. (C) Expanded view of the outer segment layer in retinal cross sections of P21 WT and *Wasf3* ^*-/-*^ mice stained with fluorophore-conjugated PNA. (D) Western blot for three representative outer segment proteins from whole retinal lysates of WT and *Wasf3* ^*-/-*^ mice at P21. Three mice of each genotype were analyzed. Each lane was loaded with 20 μg total protein. HSC70 serves as a loading control. Scale bars in all panels are 10 μm.

### *Wasf3* ^-/-^ photoreceptors produce disorganized ciliary membranes and undergo progressive degeneration

The immunostaining of outer segment proteins in *Wasf3* ^*-/-*^ mice further revealed a reduced size and dysmorphic structure of the outer segment layer (Fig. 5A,B). A particularly good example is the immunostaining of ROM1 shown in high magnification in Fig. 5B. The structural integrity of cone outer segments in *Wasf3* ^*-/-*^ mice was disrupted as well, as evident from a distorted pattern of staining with the cone outer segment marker peanut agglutinin (PNA) (Fig. 5C). The fact that both rods and cones were affected in these young animals is consistent with the finding that mutation in WASF3 in humans is linked to a condition simultaneously affecting cones and rods (36). The reduction in the size of outer segments in *Wasf3* ^*-/-*^ mice was concomitant with a reduction in the retinal contents of outer segment proteins analyzed at the same age (Fig. 5D).

A closer analysis of *Wasf3* ^*-/-*^ photoreceptors using transmission electron microscopy (TEM) displayed a striking phenotype in which large layered membranes emanate from the photoreceptor cilium instead of outer segments (Fig. 6). These membrane layers develop as early as P12. By P21, they uncontrollably expand often wrapping into large membrane whorls and persist thereafter. These data show that WASF3 plays an essential role in forming the orderly outer segment structure.

**Figure 6.**
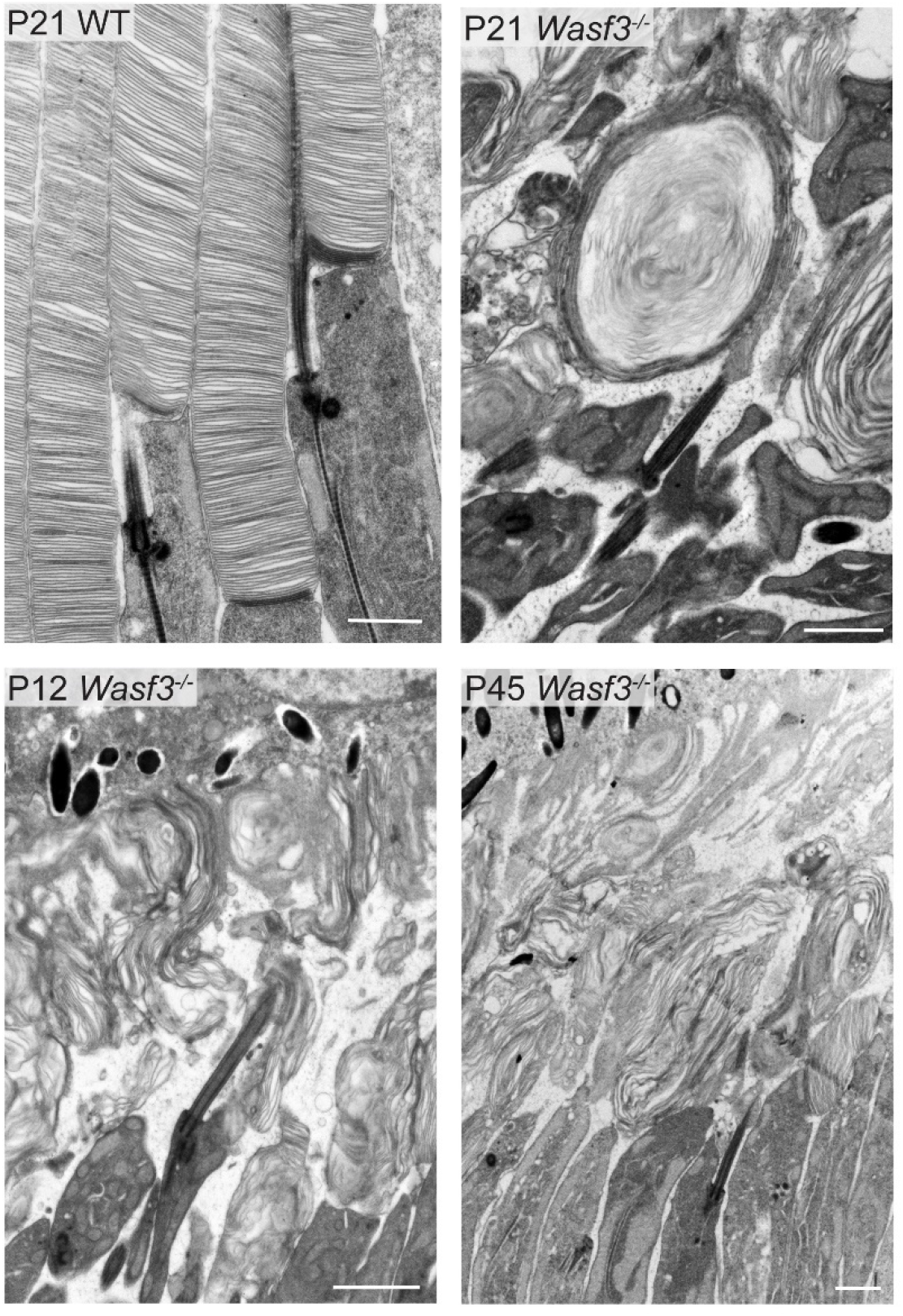
*Wasf3* ^*-/-*^ mice form disorganized ciliary membrane layers rather than outer segments. Electron micrographs of photoreceptor outer segments from WT mice or ciliary membrane layers from *Wasf3* ^*-/-*^ mice at indicated ages. Retinal sections were contrasted with tannic acid to discern disc membranes exposed to the extracellular space (darkly stained membranes) from those enclosed within the cell (lightly stained membranes). Scale bars are 1 μm.

The inability to build outer segments appears to be the only major morphological defect in young *Wasf3* ^*-/-*^ mice. This is best appreciated in thin plastic retinal sections obtained from 3-week-old mice imaged by bright field microscopy (Fig. 7A and Supplementary Fig. 2). However, photoreceptors of older *Wasf3* ^*-/-*^ mice undergo progressive degeneration, with ∼40% of cells lost at P45 (Fig. 7B,C). By P200, ∼80% of cells are lost and the few remaining photoreceptors do not possess discernable membrane structures within the outer segment layer (Fig. 7D,E). It is well-established that defects in outer segment structure often lead to the death of photoreceptor cells (1,37,38). Therefore, we reason that the disruption of the outer segment is likely to be the primary cause of photoreceptor degeneration in *Wasf3* ^*-/-*^ mice.

**Figure 7.**
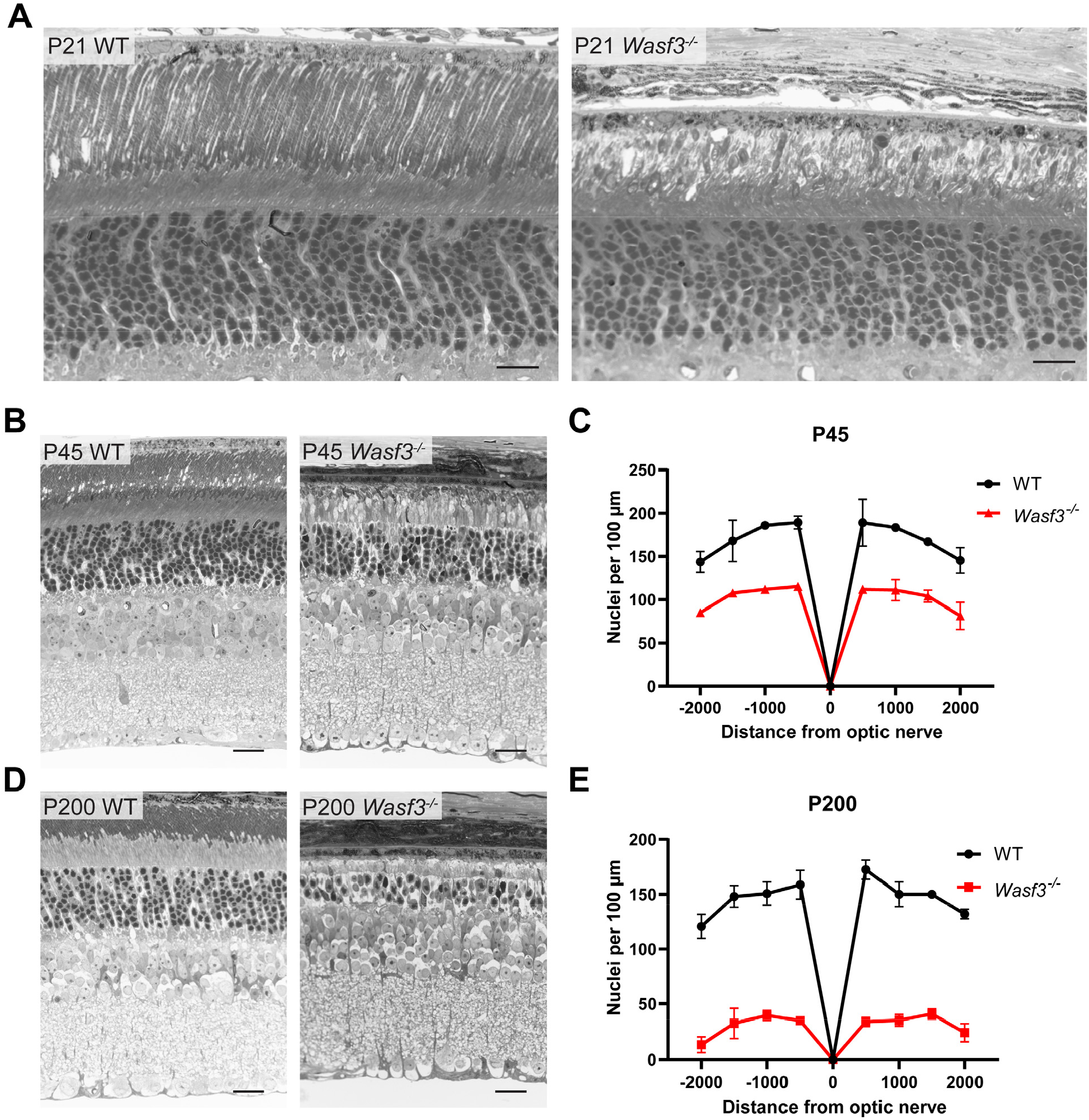
Photoreceptor cells of *Wasf3* ^*-/-*^ mice progressively degenerate. Thin plastic-embedded cross sections of WT and *Wasf3* ^*-/-*^ mouse retinas stained by Toluidine blue and imaged by light microscopy. Mice were analyzed at P21 (A), P45 (B) and P200 (D). Scale bars are 20 μm. The quantification of photoreceptor degeneration is represented by spidergram graphs showing the number of photoreceptor nuclei counted at P45 (C) and P200 (E) from eight 100-μm-wide segments of the section taken at 500 μm intervals from the optic nerve.

### Photoreceptor ciliary membranes have an intrinsic ability to form lamellar structures in the absence of actin polymerization

The photoreceptor phenotype of *Wasf3* ^*-/-*^ mice is strikingly similar to that of the rod-specific knockout of Arp2/3, in which the photoreceptor cilium elaborates extended membrane layers occasionally wrapping into whorls instead of forming an outer segment (8). In that study, we interpreted this phenotype as follows: *Because the Arp2/3 knockout is conditional, rods produce normal discs for a few weeks until their Arp2/3 content is depleted. Once Arp2/3 is gone, disc formation is halted because filamentous actin cannot be expanded to protrude the ciliary membrane. At this stage, all membrane material delivered to the outer segment can only be incorporated into the several nascent discs forming just prior to Arp2/3 depletion. Consequentially, membranes of these several nascent discs undergo uncontrolled elongation forming extended layers and whorls*. However, this interpretation does not apply to *Wasf3* ^*-/-*^ photoreceptors, because this knockout is global and WASF3 is never expressed in rods and cones of these mice. This begs the question: how could flattened membrane layers form at the cilium without actin polymerization? We envision two explanations for this phenomenon. The first is that the actin filaments are actually formed due to the activity of another nucleation promoting factor (such as WASF1), but the amount of these filaments is too small to be detected by immunofluorescence and can only produce a handful of membrane evaginations. Alternatively, the membrane material delivered to the photoreceptor cilium has an intrinsic ability to organize into lamellar sheets independent of actin polymerization.

To distinguish between these two possibilities, we analyzed the phenotype of a conditional knockout mouse in which Arp2/3 was inactivated around embryonic day 9, long before photoreceptor cells are formed. This was accomplished by breeding a mouse containing a floxed *ArpC3* allele (39) with a mouse expressing Cre recombinase behind the Six3 promoter (40). Unlike nucleation promoting factors, there is no redundancy to Arp2/3 function, which means that the Six3Cre/*ArpC3* conditional knockout mouse completely prevents branched actin polymerization in knockout photoreceptors. We found that this early Arp2/3 knockout severely disrupts retinal development, including proper formation of retinal layers (Fig. 8A and Supplementary Fig. 3). However, photoreceptors were still formed in these retinas and produced whorls of membrane layers emanating from their cilia (Fig. 8B). This result argues that the ability to form layered structures is an intrinsic property of photoreceptor ciliary membranes that does not require priming by actin polymerization (potential mechanisms are considered in the Discussion). However, this spontaneous formation of membrane layers is not sufficient to build an orderly outer segment structure.

**Figure 8.**
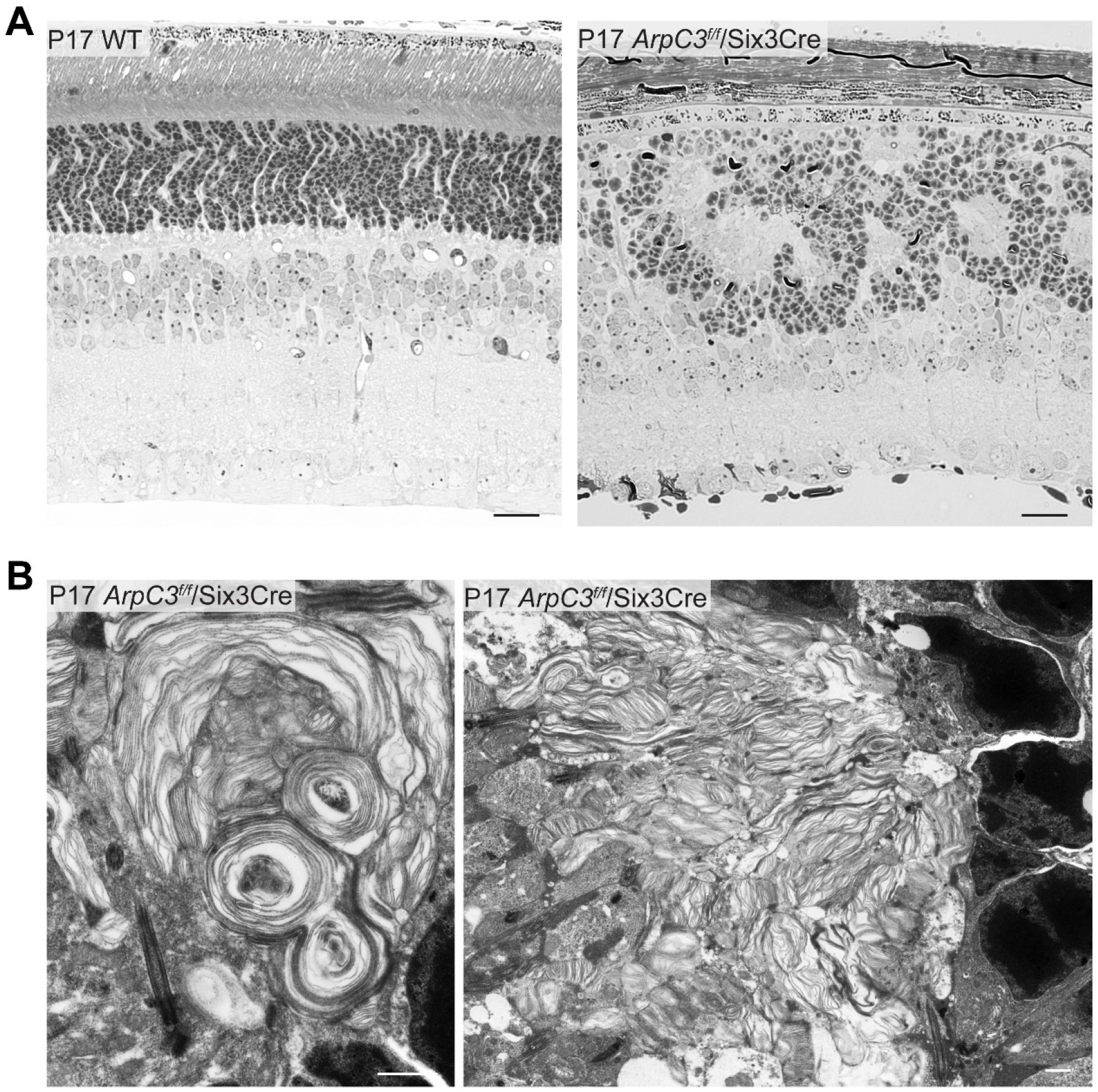
The early knockout of Arp2/3 disrupts retinal development while preserving the ability of photoreceptors to produce ciliary membrane layers. (A) Thin plastic-embedded cross sections of P17 WT and Six3Cre/*ArpC3* ^*-/-*^ mouse retinas stained by Toluidine blue and imaged by light microscopy. Scale bars are 20 μm. (B) Electron micrographs of photoreceptor ciliary membrane layers from P17 Six3Cre/*ArpC3* ^*-/-*^ mice. Scale bars are 1 μm.

Although beyond the scope of this study, another interesting observation from the Six3Cre/*ArpC3* conditional knockout mouse is that the role of branched actin in retinal development is not limited to disc morphogenesis. The eye diameter of these animals is reduced and the laminar organization of their retinas is disrupted. The latter phenotype resembles some previously described models in which the apical-basal polarity of the neuroepithelium is disrupted (41-43).

## Discussion

The central observation of this study is that the pentameric WAVE complex is an essential component of the photoreceptor disc morphogenesis machinery. This complex acts as the nucleation promoting factor for Arp2/3, which drives actin polymerization at the outer segment base to evaginate the ciliary membrane and initiate the formation of a new disc. We also identified specific protein isoforms representing each subunit of this WAVE complex, including the indispensable WASF3 isoform.

The role of WASF3 in outer segment formation is particularly intriguing because the WASF3 knockout mouse has no other overt morphological or behavioral phenotypes (25). Furthermore, cone-rod dystrophy was the only abnormality described in a human patient having a splice site mutation in the *WASF3* gene (36). Thus, driving actin polymerization during photoreceptor disc morphogenesis is the only indispensable function of WASF3 in normal cells identified to date. Outside of normal cell functions, WASF3 upregulation in certain types of cancer cells was shown to promote metastasis through enhanced actin polymerization (30). Since targeting WASF3 has been suggested as a potential cancer therapy (21,44), special attention to photoreceptor cell health should be considered during the testing of such a treatment.

One other molecular player reported to interact with WASF3 is PCARE, a protein analyzed in the study detailed above, in which the authors proposed that PCARE recruits WASF3 to the site of disc morphogenesis to initiate actin network expansion (19). However, we did not detect PCARE in either WASF3 or ABI1 immunoprecipitates from outer segment lysates, despite robustly identifying other components of the WAVE complex in both cases. Of note, our experiments were performed with purified rod outer segments, whereas their experiments were conducted in cell culture. One potential explanation for this discrepancy is that the affinity between these proteins is rather weak and preservation of the complex requires protein overexpression. It is also possible that these proteins do not interact directly but rather through one of many other components of the actin cytoskeleton found in PCARE precipitates from cell culture (19), including actin itself shown to interact with each of these proteins (17,19). Regardless, the specific role of PCARE in regulating the actin network at the disc morphogenesis site requires further investigation.

Another major impact of the present study is that it allows discernment of which aspects of photoreceptor disc formation are dependent on actin polymerization and which are not. The cycles of actin polymerization and depolymerization initiate the formation of each new disc in a periodic fashion, thereby assuring that the membrane material delivered to the outer segment is evenly allocated into an orderly stack of discs. On the other hand, our data show that photoreceptor ciliary membranes are able to form flattened layers without actin polymerization. This ability is not trivial because no other membrane expansions of the primary cilium build layered structures (e.g. (19,45-48)). Therefore, photoreceptor ciliary membranes must contain specific molecular players that promote the membrane bending and cross-membrane adhesion required for these layers to be formed.

A number of proteins have been implicated in performing membrane bending in the outer segment. They are illustrated schematically in Fig. 9. The best studied is the oligomeric complex between peripherin-2 and ROM1, which bends disc membranes in the inward direction towards the cytoplasm and supports membrane curvature along the entire circumference of mature disc rims (37,49,50). Membrane bending in the opposite direction, away from the cytoplasm, is thought to be facilitated by prominin, a protein involved in forming various protrusions of the plasma membrane in other cells (51). Prominin is located at the expanding edges of newly forming photoreceptor discs and, therefore, may stabilize the membrane bend at this location (52). It is likely that, in the absence of actin polymerization, the same proteins are responsible for the formation of the membrane bends present in the layers of photoreceptor ciliary extensions.

**Figure 9.**
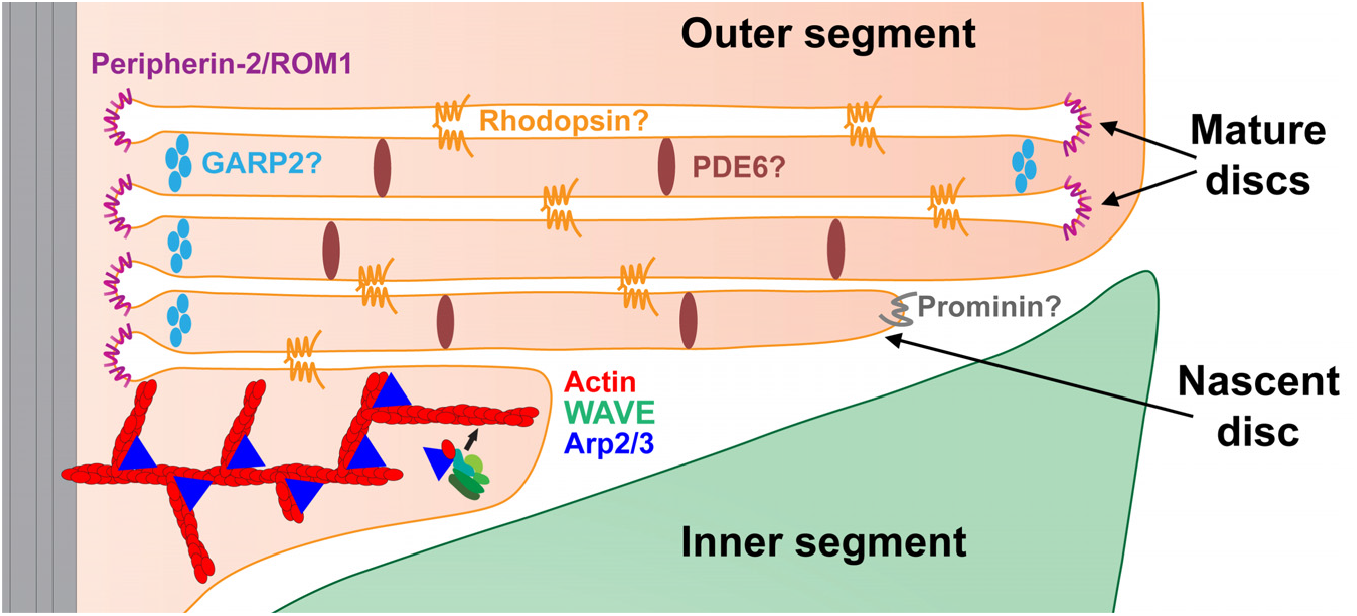
Cartoon illustrating the role of the WAVE complex in initiating photoreceptor disc formation and functions of proteins implicated in membrane bending and trans-membrane adhesion. The WAVE complex cooperates with Arp2/3 to drive actin polymerization evaginating each nascent disc. A more detailed illustration of WAVE-Arp2/3-driven actin polymerization is presented in Fig. 1. Extended oligomeric complexes of peripherin-2/ROM1 support the highly curved edges along the entire circumference of mature discs. The leading edges of expanding nascent discs are enriched with prominin, which is thought to promote membrane bending at this location. Homotypic rhodopsin interactions have been hypothesized to provide membrane-to-membrane adhesion across the extracellular space between nascent discs and the intradiscal space of mature discs. PDE6 and/or GARP2 have been hypothesized to connect discs across the cytoplasm throughout disc surface and near the disc rim, respectively. Proteins whose role in membrane bending or adhesion remains hypothetical are denoted with a question mark. See Discussion for details.

The stabilization of flattened membrane layers requires adhesive interactions on both intracellular and extracellular surfaces of each apposing membrane. Little is known about the proteins that fulfill these roles, although several candidates have been proposed (Fig. 9). It was hypothesized that, on the extracellular surface (eventually turning into the intradiscal space of mature discs as they enclose), membrane-to-membrane adhesion is achieved by multiple weak interactions between rhodopsin molecules across the extracellular/intradiscal space (53). This idea is generally consistent with the fact that the membrane extensions forming at the cilium of rhodopsin knockout mice do not organize into membrane layers (54). The presence of linkers connecting disc membranes across the cytoplasmic space has been documented in multiple ultrastructural studies (see (50) for a recent update). Two proteins, PDE6 and GARP2, were hypothesized to play a role in forming these links (50), although neither of them has been validated in direct experiments.

Another aspect of disc morphogenesis that occurs independently of actin polymerization is disc enclosure. The tissue processing technique employed in our TEM experiments darkly stains membranes exposed to the extracellular space and lightly stains those enclosed inside the outer segment (5). The large membrane whorls emanating from *Wasf3* ^*-/-*^ and Six3Cre/*ArpC3* photoreceptor cilia contain both lightly stained and darkly stained membranes (e.g. Figs. 6 and 8B), indicating that some of these membrane layers can still be enclosed despite their malformation in the absence of F-actin. This conclusion is consistent with our previous observation that neither cytochalasin D treatment nor a rod-specific Arp2/3 knockout prevented disc membrane enclosure (8).

In conclusion, our demonstration that the WAVE complex is an indispensable component of the disc formation machinery brings us one step closer to elucidating the entire mechanism of photoreceptor outer segment morphogenesis. The immediate goals of future studies are to identify the molecular players acting upstream of the WAVE complex and to elucidate whether the specialization of this complex in disc morphogenesis is conferred exclusively by WASF3 or whether other subunits of the WAVE complex contribute as well.

## Methods

### Animals

Mice handling was performed in accordance with the approved protocol by the Institutional Animal Care and Use Committees of Duke University (A240-19-11). WT C57BL/6J mice (stock #000664) were obtained from Jackson Labs (Bar Harbor, ME). *ArpC3*^*f/f*^ mice (39) were kindly provided by Dr. Scott Soderling (Duke University) and crossed with Six3-Cre mice (40) obtained from Jackson Labs (stock #019755) to generate *ArpC3*^*f/f*^/Six3-Cre mice. *Wasf3*^*-/-*^ mice (22) were generously provided by Dr. John Cowell (Georgia Cancer Center). All mice were housed under a 12/12 hour diurnal light cycle.

### Antibodies

We used rabbit anti-WASF3 (#2806; Cell Signaling Technology) and mouse anti-ABI1 (D147-3; MBL) for immunoprecipitation experiments. Mouse monoclonal antibody 4D2 against the N-terminus of rhodopsin and mouse anti-CNGα1 were kindly gifted to us from Robert S. Molday (University of British Columbia, Vancouver, Canada) and used for western blotting and/or immunofluorescence staining. Rabbit polyclonal anti-peripherin-2 antibody was gifted by Gabriel H. Travis (University of California, Los Angeles) and used for western blotting and/or immunofluorescence staining. We also used the following commercial antibodies for western blotting and/or immunofluorescence staining: rabbit anti-WASF3 (#2806; Cell Signaling Technology), mouse anti-ARP3 (ab49671; Abcam), rabbit anti-CYFIP2 (ab95969; Abcam), rabbit anti-CYFIP1 (#PA5-31984; Thermo Fisher Scientific), rabbit anti-NCKAP1 (12140-1-AP; Proteintech), mouse anti-BRK1 (SC-390459; Santa Cruz), rabbit anti-ABI1 (#39444; Cell Signaling Technology), mouse anti-GRK1 (MA1-720; Thermo Fisher Scientific), anti-ABCA4 antibody (SC-21460; Santa Cruz) and rabbit anti-HSC70 (ADI-SPA-819; Enzo Life Sciences). We generated a custom polyclonal sheep anti-ROM1 antibody and used it for western blotting and immunofluorescence staining. This ROM1 antibody was generated against the peptide CIDGEGEAQGYLFPAGLKDM representing a C-terminal bovine ROM1 sequence using methods described previously (55). The antibody was affinity-purified using the corresponding peptide attached to SulfoLink beads (ThermoFisher Scientific, Waltham, MA) via the N-terminal cysteine residue.

### Immunofluorescence

Mice were deeply anesthetized and transcardially perfused using fixative solution containing 4% paraformaldehyde in 80 mM PIPES (pH6.8), 5 mM EGTA, and 2mM MgCl_2_, resulting in exsanguination. The eyes were carefully removed and post-fixed in the same fixative for two hours at 22°C then washed three times with PBS. The eyes were then dissected into eyecups and embedded in 2.5% agarose (SKU VF-AGT-VM; Precisionary) and cut into 100 μm thick slices by a vibratome (VT1200S; Leica). Sections were blocked in PBS containing 7% donkey serum and 0.5% Triton X-100 for 1 h at 22°C. The sections were then incubated overnight with primary antibodies. After three washes, the sections were stained for 2 h at 22°C with appropriate Alexa Fluor conjugated secondary antibodies (Invitrogen), 10 μg/mL Hoechst (H3569; Thermo Fisher Scientific), 1 μg/mL WGA conjugated with Alexa Fluor 594 (W11262; Thermo Fisher Scientific), 1 μg/mL PNA conjugated with Alexa Fluor 647 (L32460; Thermo Fisher Scientific) and/or 130 nM phalloidin conjugated with Alexa Fluor 488 (A12379; Thermo Fisher Scientific). The sections were washed three times in PBS and mounted onto slides with Shandon Immu-Mount (9990402; Thermo Fisher Scientific) and coverslipped. The sections were imaged with a confocal microscope (Eclipse 90i and A1 confocal scanner; Nikon) with a 60× objective (1.49 NA Plan Apochromat VC; Nikon) using Nikon NIS-Elements software. Image processing included adjusting brightness and contrast and was performed with ImageJ and/or Nikon NIS-Elements software.

### Electron Microscopy

Fixation and processing of mouse eyes for thin plastic sections was performed as described previously (5). Mice were deeply anesthetized and transcardially perfused with 2% paraformaldehyde, 2% glutaraldehyde, and 0.05% calcium chloride in 50 mM MOPS (pH 7.4) resulting in exsanguination. The eyes were enucleated and fixed for an additional 2 h in the same buffer at 22°C. To obtain thin plastic retinal sections, eyecups were cut in half through the optic nerve and embedded in Spurr’s resin (Electron Microscopy Sciences). The embedded retinal cross sections were cut through the optic nerve in 500 nm slices and stained with methylene blue for light microscopy as in (56).

For electron microscopy, eyes were fixed and processed as described above for thin plastic sections. The fixed eyes were washed in PBS, eyecups were dissected and embedded in PBS containing 2.5% agarose (KU VF-AGT-VM; Precisionary), and cut into 200 μm thick slices on a Vibratome (VT1200S; Leica) (5). The Vibratome sections were stained with 1% tannic acid (Electron Microscopy Sciences) and 1% uranyl acetate (Electron Microscopy Sciences), gradually dehydrated with ethanol and embedded in Spurr’s resin. 70 nm sections were cut, placed on copper grids, and counterstained with 2% uranyl acetate and 3.5% lead citrate (19314; Ted Pella). The samples were imaged on a JEM-1400 electron microscope (JEOL) at 60 kV with a digital camera (Biosprint 16; AMT).

### Rod Outer Segment Purification

Osmotically intact mouse rod outer segments (ROS) were isolated as described previously (57) with small modifications. Retinas from at least four 2-month old mice were isolated in mouse Ringer’s solution containing 130 mM NaCl, 3.6 mM KCl, 2.4 mM MgCl_2_, 1.2 mM CaCl_2_, 10 mM Hepes (pH 7.4) adjusted to 314 mosM. The retinas were pooled into 400 μl ice-cold Optiprep (8%; Sigma-Aldrich) in Ringer’s solution and vortexed at maximum speed for 60 s in a 1.5 ml Eppendorf tube. The tube was then centrifuged at 200×g for 30 s to sediment large retinal debris. 350 μl of the supernatant was loaded on top of a 1.8 ml step gradient composed of 10 and 18% OptiPrep in Ringer’s solution and centrifuged for 30 min at 20,000 rpm in a swing bucket SW-55 rotor (Beckman Coulter) at 4°C. ROS were carefully collected from the 10/18% OptiPrep interface, diluted in Ringer’s solution, centrifuged at 10,000×g and the resulting pellet was washed once with Ringer’s solution before lysing in solubilization buffer.

Rod outer segment purification from bovine retinas was performed following a modified procedure from (58), as described previously (59). Briefly, 100 frozen bovine retinas (T.A. & W.L. Lawson Co) were thawed and resuspended in 200 ml of buffer A (20 mM Hepes, 100 mM KCl, 2 mM MgCl2, and 0.1 mM EDTA; pH 7.4) containing 25% sucrose. Outer segments were detached from the retinas by swirling in a 1 liter Erlenmeyer flask and separated from retinal debris by centrifugation at 3700×g for 6 min. The supernatant was diluted with an equal volume of buffer A, and ROSs were pelleted by centrifugation at 6300×g for 8 min. The pellet was resuspended in buffer A containing 20% sucrose, applied on a 27/32% step gradient of sucrose and centrifuged for 1 h in an SW-28 rotor at 83,000×g. ROSs were collected from the 27/32% sucrose interphase, centrifuged at 10,000×g and the resulting pellet was washed once with buffer A before lysing in solubilization buffer.

### Mass Spectrometry

Briefly, proteins solubilized in 2% SDS, 100 mM Tris·HCl (pH 8.0) were reduced with 10 mM DTT (D0632; Sigma-Aldrich), alkylated with 25 mM iodoacetamide (I1149; Sigma-Aldrich) and subjected to tryptic hydrolysis using the HILIC beads SP3 protocol (60). The resulting peptides were analyzed with a nanoAcquity UPLC system (Waters) coupled to an Orbitrap Q Exactive HF mass spectrometer (Thermo Fisher Scientific) employing the LC-MS/MS protocol in a data-independent acquisition mode. The peptides were separated on a 75 μm × 150 mm 1.7 μm C18 BEH column (Waters) using a 90 min gradient of 6% to 30% of acetonitrile in 0.1% formic acid at a flow rate of 0.3 ml/min at 45°C. Eluting peptides were sprayed into the ion source of the Orbitrap Q Exactive HF at a voltage of 2.0 kV and data acquisition was performed for top 15 precursors, automatic gain control was e^6^ and injection time -100 msec. Progenesis QI Proteomics software (Waters) was used to assign peptides to the features and generate searchable files which were submitted to Mascot (version 2.5) for peptide identification. For peptide identification we searched against the UniProt reviewed mouse database (September 2019 release) using carbamidomethyl at Cys as a fixed modification and Met oxidation as a variable modification. Only nonconflicting peptides were used for protein identification and quantification and only proteins with two or more peptides and protein confidence P<0.05 were selected as confidently identified.

### Western Blotting and Immunoprecipitation

To compare retinal protein abundance between WT and *Wasf3*^*-/-*^ mice, the retinas were sonicated in 200 μL of 2% SDS containing protease inhibitor mixture (eComplete; Roche) in PBS. Next, the lysates were centrifuged at 20,000 × g for 10 min at 22 °C, and the resulting supernatants were normalized by total protein concentration measured by the RC DC Protein Assay kit (Bio-Rad). Equal volumes of retinal lysate containing 10 μg total protein from each mouse genotype were subjected to SDS/PAGE. Proteins were transferred to low-fluorescence PVDF membrane (Bio-Rad), blocked with Intercept Blocking Buffer (927-70001; LiCor Biosciences) and probed with primary antibodies listed above and goat secondary antibodies conjugated with Alexa Fluor 680 or 800 (Invitrogen) for detection on an Odyssey infrared imaging system (LiCor Bioscience).

Lysates for immunoprecipitation were produced by gently solubilizing purified outer segments in buffer containing 50 mM Tris, 150 mM NaCl, 1% NP-40, 10% glycerol, pH 7.5 and protease inhibitor mixture (eComplete; Roche) on ice. The lysate was cleared by centrifugation at 10,000×g for 20 minutes at 4 °C. The lysate was split between two low retention microcentrifuge tubes (02-681-311; Fisher Scientific) containing experimental (anti-WASF3 or anti-ABI1) or control (anti-IgG) antibodies and incubated overnight at 4 °C. The antibodies and any bound proteins were purified by incubating the lysates with protein A/G magnetic beads (Thermo Scientific), washing the beads three times with lysis buffer and eluting bound proteins with 2% SDS in PBS.

## Acknowledgments

This work was supported by the National Institutes of Health grants EY012859 (V.Y.A.), EY005722 (V.Y.A.), the Career-Starter Research Grant from the Knights Templar Eye Foundation (W.J.S.) and an Unrestricted Award from Research to Prevent Blindness Inc. (Duke University).

## Author contributions

W.J.S. and V.Y.A. conceived and designed the experiments. W.J.S., N.F.S. and N.P.S. performed the experiments. W.J.S., N.F.S. and N.P.S. and V.Y.A. analyzed the data. W.J.S. and V.Y.A. wrote the manuscript. All authors edited the manuscript.

## Supplementary Information

**Supplementary Figure 1.**
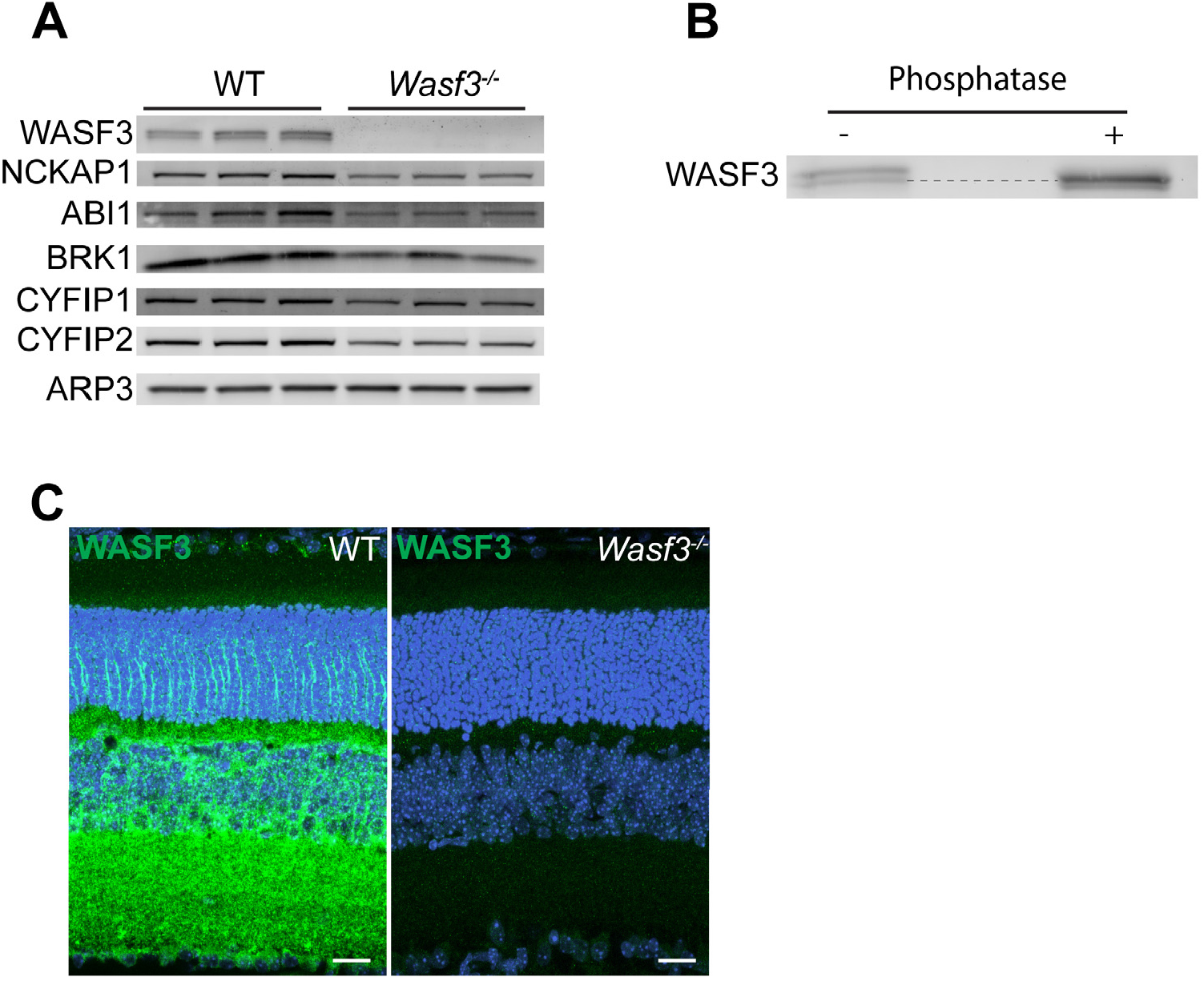
Confirmation of the loss of WASF3 in the retina of *Wasf3* ^*-/-*^ mice. (A) Western blots of WAVE complex subunits from retinal lysates of WT and *Wasf3* ^*-/-*^ mice at P21. Three mice of each genotype were analyzed. Each lane was loaded with 20 μg total protein. (B) Retinal cross sections of WT and *Wasf3* ^*-/-*^ mice at P21 immunostained with anti-WASF3 antibody (green). Nuclei are stained with Hoechst (blue). The scale bars are 20 μm. (C) Western blot of WASF3 from WT retinal lysate, which was either treated or not treated with calf intestinal phosphatase. The horizontal dashed line highlights the location of the lower band.

**Supplementary Figure 2.**
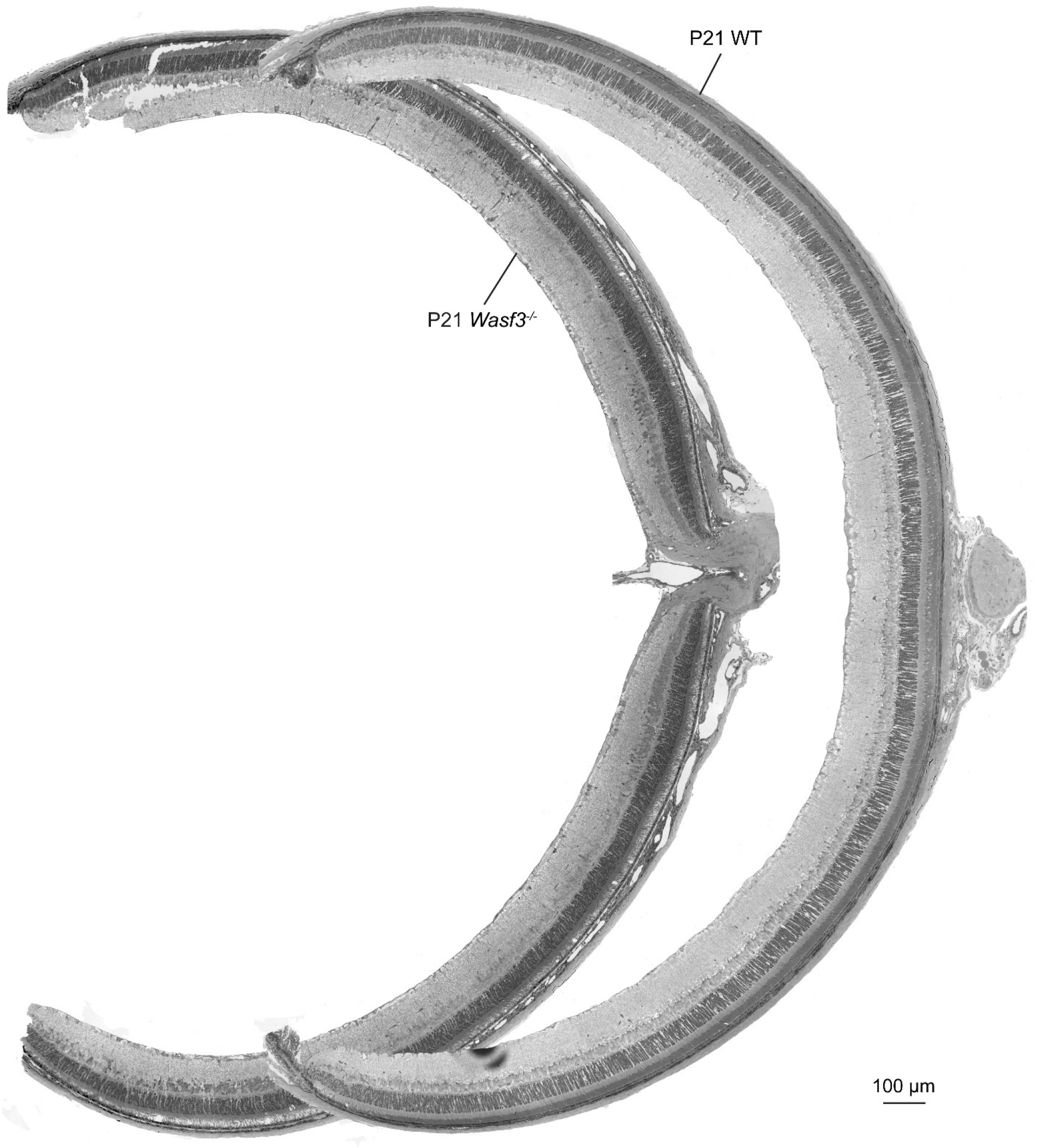
Representative tile scanned images of the entire retina from P21 WT and *Wasf3* ^*-/-*^ mice.

**Supplementary Figure 3.**
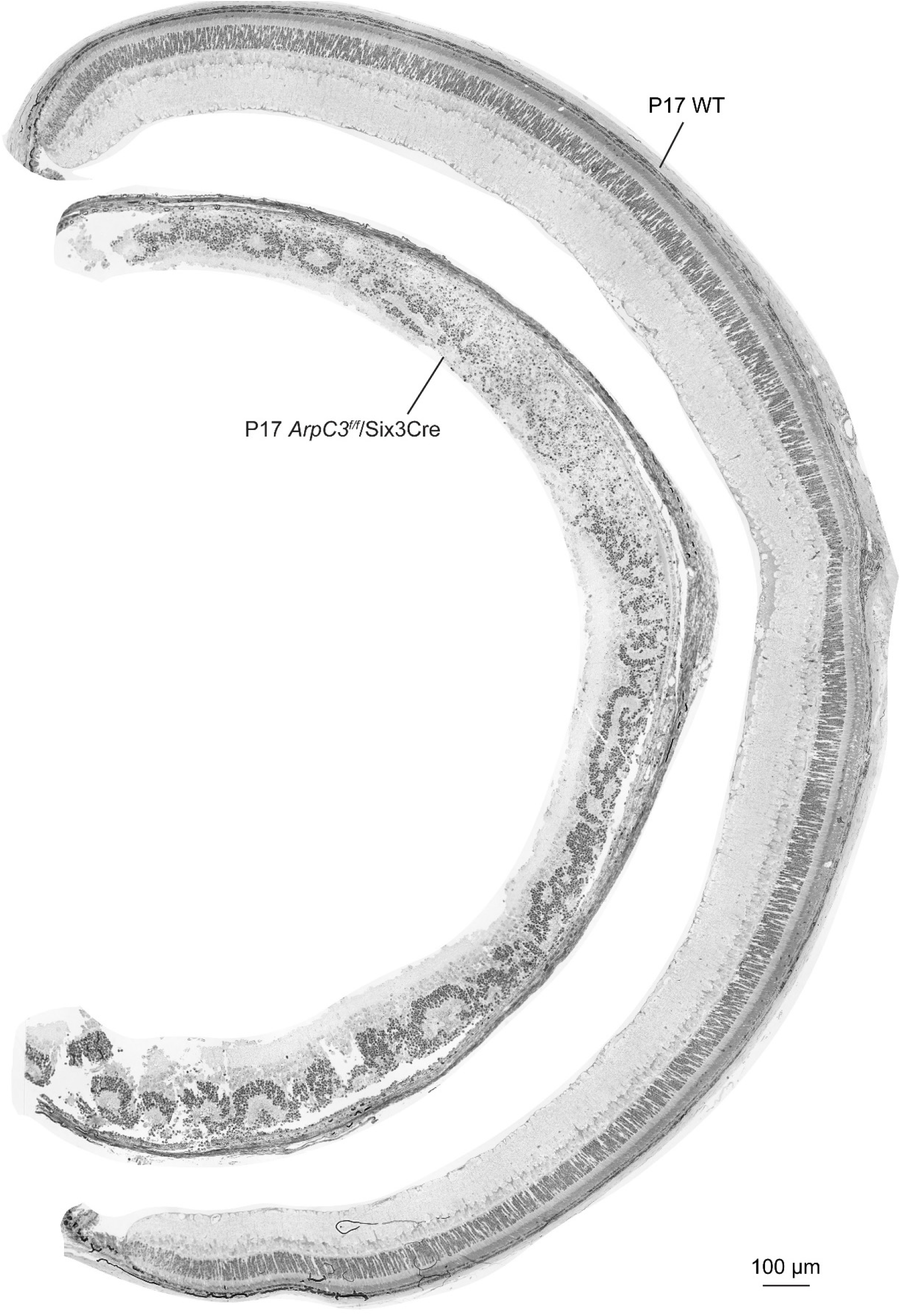
Representative tile scanned images of the entire retina from P17 WT and *Six3Cre/ArpC3* ^*-/-*^ mice.

**Supplementary Figure 4.**
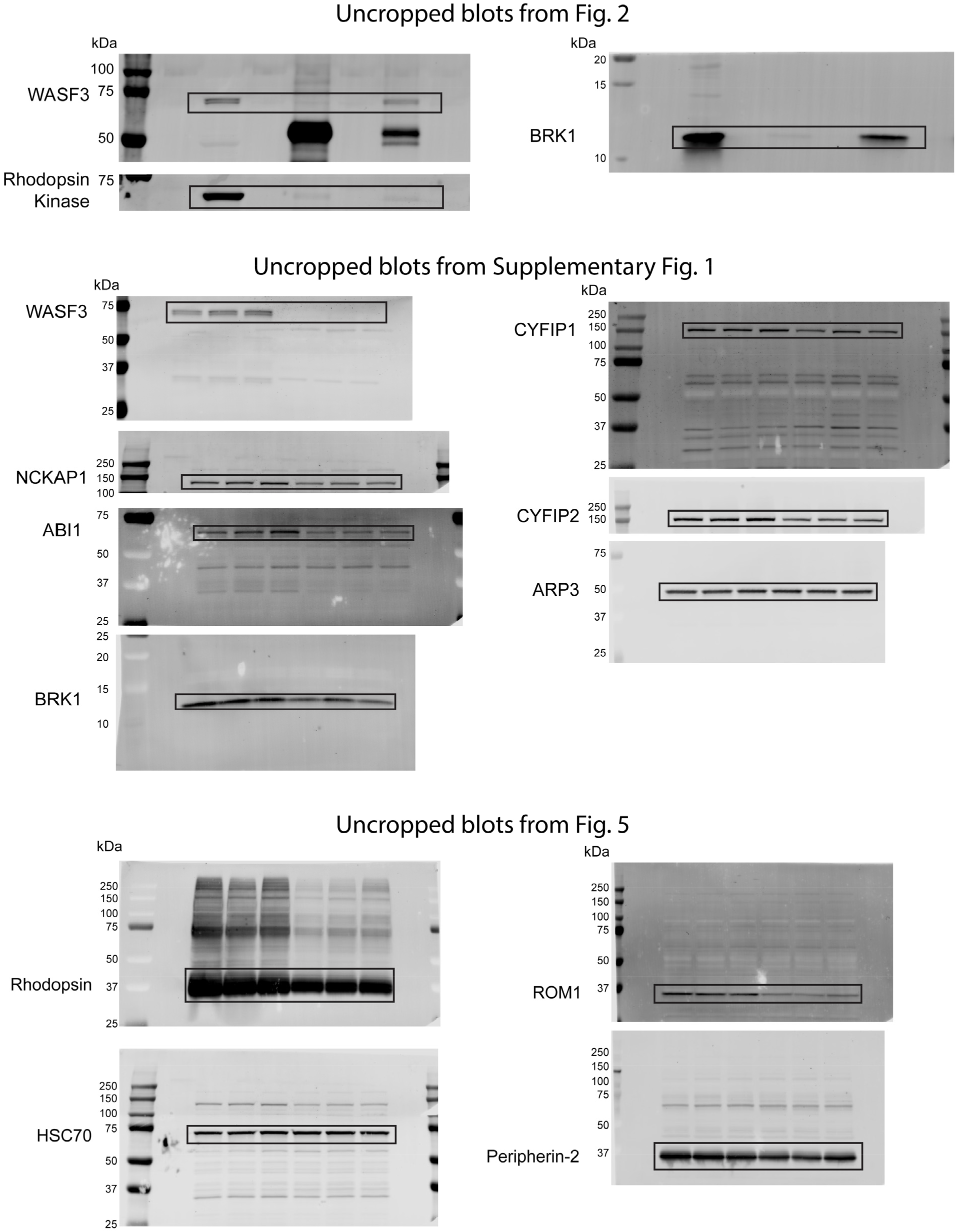
The uncropped images of western blot in this study. The black boxes indicate the cropped region shown in the main figures.

**Supplementary Table 1**. Proteins confidently identified by mass spectrometry in a preparation of isolated bovine rod outer segments. Shown are proteins represented by two or more identified peptides with protein confidence value <0.05. The data are sorted by the number of spectral counts, except for the three WAVE complex subunits shown at the top and highlighted in yellow.

**Supplementary Table 2**. Proteins confidently identified by mass spectrometry in immunoprecipitates from lysates of purified mouse rod outer segments obtained with anti-WASF3 or control IgG antibodies. For each protein, the total number of unique peptides, the confidence score associated with the protein’s identification, and the total ion intensity of all identified peptides are shown. Protein enrichment was calculated by dividing the total ion intensity of the protein’s identified peptides in the WASF3 immunoprecipitate by the protein’s total ion intensity in the control immunoprecipitate. Both average and median enrichment values across four independently conducted experiments are listed. Proteins in the table are sorted from highest median enrichment to lowest. The WAVE complex subunits are highlighted in yellow. Proteins included in the table are: 1) identified in at least three experiments; 2) have at least 2 identified peptides in any given experiment; 3) enriched, on average, by at least 2-fold across all experiments.

**Supplementary Table 3**. Proteins confidently identified by mass spectrometry in immunoprecipitates from lysates of purified mouse rod outer segments obtained with anti-ABI1 or control IgG antibodies. For each protein, the total number of unique peptides, the confidence score associated with the protein’s identification, and the total ion intensity of all identified peptides are shown. Protein enrichment was calculated by dividing the total ion intensity of the protein’s identified peptides in the WASF3 immunoprecipitate by the protein’s total ion intensity in the control immunoprecipitate. Average enrichment values between two independently conducted experiments are listed. Proteins in the table are sorted from highest to lowest enrichment. The WAVE complex subunits are highlighted in yellow. Proteins included in the table are: 1) identified in both experiments; 2) have at least 2 identified peptides in at least one experiment; 3) enriched by at least 2-fold after averaging the values from both experiments.

**Supplementary Movie 1. A video showing a 3D volume representation of a Z-stack confocal image of a P21 WT retinal cross section stained with phalloidin to label F-actin**. The video zooms in on the inner-outer segment juncture.

**Supplementary Movie 2. A video showing a 3D volume representation of a Z-stack confocal image of a P21 *Wasf3*** ^***-/-***^ **retinal cross section stained with phalloidin to label F-actin**. The video zooms in on the inner-outer segment juncture.

